# Improvement of Attention-Deficit/Hyperactivity Disorder (ADHD) Symptoms in School-Aged Children, Adolescents, and Young Adults With Autism via a Digital Smartglasses-Based Socioemotional Coaching Aid: An Efficacy Study

**DOI:** 10.1101/165514

**Authors:** Arshya Vahabzadeh, Neha U. Keshav, Joseph P. Salisbury, Ned T. Sahin

## Abstract

**Background:** People with autism spectrum disorder (ASD) commonly experience symptoms related to attention-deficit/hyperactivity disorder (ADHD), including hyperactivity, inattention, and impulsivity. One-third of ASD cases may be complicated by the presence of ADHD. Individuals with dual diagnoses face greater barriers to accessing treatment for ADHD and respond less positively to primary pharmacologic interventions. Nonpharmacologic technology-aided tools for hyperactivity and inattention in people with ASD are being developed, although research into their efficacy and safety remains limited.

**Objective:** The objective of this preliminary study was to describe the changes in ADHD-related symptoms in children, adolescents, and young adults with ASD after use of the Empowered Brain system, a behavioral and social communication aid for ASD running on augmented reality smartglasses.

**Methods:** We recruited 8 children, adolescents, and young adults with ASD (male to female ratio of 7:1, mean age 15 years, range 11.7-20.5 years) through a Web-based research signup form. The baseline score on the hyperactivity subscale of the Aberrant Behavioral Checklist (ABC-H), a measure of hyperactivity, inattention, and impulsivity, determined their classification into a high ADHD-related symptom group (n=4, ABC-H ≥13) and a low ADHD-related symptom group (n=4, ABC-H <13). All participants received an intervention with Empowered Brain, where they used smartglasses-based social communication and behavioral modules while interacting with their caregiver. We then calculated caregiver-reported ABC-H scores at 24 and 48 hours after the session.

**Results:** All 8 participants were able to complete the intervention session. Postintervention ABC-H scores were lower for most participants at 24 hours (n=6, 75%) and for all participants at 48 hours (n=8, 100%). At 24 hours after the session, average participant ABC-H scores decreased by 54.9% in the high ADHD symptom group and by 20% in the low ADHD symptom group. At 48 hours after the session, ABC-H scores compared with baseline decreased by 56.4% in the high ADHD symptom group and by 66.3% in the low ADHD symptom group.

**Conclusions:** This study provides initial evidence for the efficacy of the Empowered Brain system in reducing ADHD-related symptoms, such as hyperactivity, inattention, and impulsivity, in school-aged children, adolescents, and young adults with ASD. This digital smartglasses intervention can potentially be targeted at a broader array of mental health conditions that exhibit transdiagnostic attentional and social communication deficits, including schizophrenia and bipolar disorder. Further research is required to understand the clinical importance of these observed changes and to conduct longitudinal studies on this intervention with larger sample sizes.

## Introduction

Autism spectrum disorder (ASD) is a lifelong developmental disorder characterized by challenges in social communication and the presence of repetitive behaviors or restricted interests. Many people with ASD experience symptoms of inattention and hyperactivity, and approximately one-third of people with ASD have diagnosable attention-deficit/hyperactivity disorder (ADHD) [1,2]. There are considerable gaps in knowledge in how to provide optimal assessment and management of this group of patients [3]. Early diagnosis and intervention are beneficial for both ASD [4] and ADHD [5]. Evidence from genetic, cognitive, and behavioral research suggests that when ADHD and ASD co-occur, they may be considered a separate overarching condition [6-8], and a variety of ASD-ADHD developmental subtypes have already been proposed [9].

The combination of ASD and ADHD has been linked to greater cognitive impairment [10,11], general psychopathology [12,13], emotional processing impairment [14], and to significantly higher rates of some hyperactivity and impulsivity symptoms than in individuals with ADHD alone [15]. Furthermore, the co-occurrence of ADHD and ASD is associated with a greater risk of developing other psychiatric disorders, such as schizophrenia, bipolar disorder, and anxiety disorder, than in controls and individuals with either condition alone [16]. The pervasive nature of impairments in social communication and attention across psychiatric disorders may suggest that these deficits should be investigated transdiagnostically, an approach advocated by the US National Institute of Mental Health Research Domain Criteria framework [17]. Attentional and social communication deficits have been identified in many disorders of brain function, including schizophrenia [18,19], bipolar disorder [20,21], anxiety disorders [22], and traumatic brain injury [23,24]. Therefore, the study of individuals with combined ASD and ADHD can help elucidate the basis of a wide variety of disorders of the brain.

It is therefore important to address ADHD. While the leading approach has been psychopharmacologic medication, people with co-occurring ASD and ADHD have been found to be less likely to receive appropriate treatment for their ADHD [15] and appear to respond less favorably to treatment than do individuals with ADHD alone [25]. Additional concerns about ADHD treatment, in particular stimulant medication, focus on their long-term effectiveness [26], side effects [27], and parental reservation about their use [28]. Yet evidence also shows that leaving individuals with untreated ADHD may lead to considerable negative social and behavioral sequelae, including greater risk of academic failure [29], alcohol and drug use [30], and contact with the criminal justice system.

There has been growing interest in the use of cognitive training in ADHD, a nonpharmacologic approach that may use neurofeedback or novel digital approaches, or both. Recent studies have shown promise [31,32], although historic interventions have raised questions regarding their effectiveness [33]. There is also concern that technology may actually prove to be distracting and reduce learners’ attentiveness to educational tasks [34-36].

Little research has described the impact of digital interventions on people with ASD who demonstrate ADHD symptoms [37]. Some preliminary research has shown the utility of augmented reality interventions in ASD samples [38-40]. Intervention based on augmented reality has also been shown to help to improve ADHD symptoms in people with ASD, with improvements in both selective and sustained attention in children with ASD [38]. We have previously described the delivery of social communication coaching on augmented reality smartglasses via Empowered Brain (previously the Brain Power Autism System;Brain Power LLC) [41]. Our early pilot report on 2 boys with ASD demonstrated short-term improvements in the hyperactivity subscale of the Aberrant Behavioral Checklist (ABC-H) [41], a validated instrument that assesses hyperactivity, impulsivity, attention, and noncompliance [42]. The ABC-H has previously been used as a key outcome measure in ADHD treatment studies in children with ASD [43-47].

## Objectives

In this pilot study, we explored the short-term effect of an Empowered Brain intervention on ADHD symptoms in a group of 8 children, adolescents, and young adults with ASD. We documented caregiver-reported ADHD symptoms as measured by the ABC-H. Further, we discuss the implications of these results on future research in the field.

### The Empowered Brain System

Empowered Brain is a combination of modern smartglasses and educational modules targeting socioemotional and behavioral management skills [41,48]. Smartglasses are lightweight head-worn computers with a small transparent display that can provide guidance to users through both visual and audio cues (Figure 1 [52]). Empowered Brain can collect a wide variety of user data through an in-built sensor array that includes a camera, microphone, touchpad, “blink” sensor, gyroscope, and accelerometer. Empowered Brain includes modules that use these sensors to deliver social communication and cognitive skills coaching. This digital approach may be particularly valuable to both people with ASD [49] and people with ADHD [50,51].

**Figure 1.**
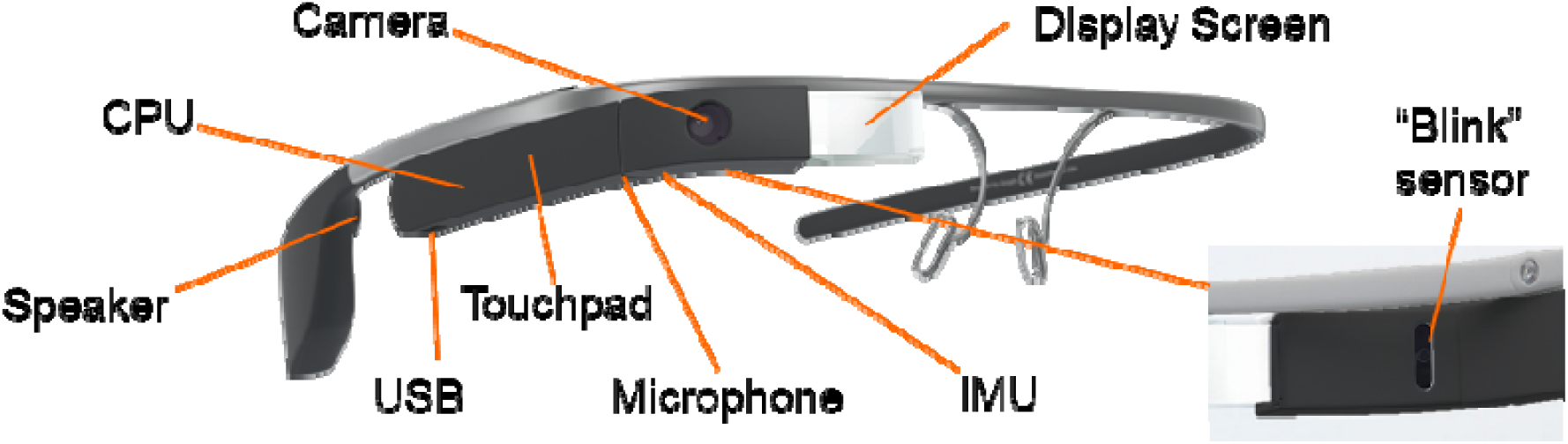
*Google Glass*, prototypical head-worn smartglasses with in-built sensors, as well as a small screen and a bone conduction speaker to provide a private audio-visual experience. Empowered Brain integrates Google Glass with a range of assistive and educational modules. User-centered design is an critical part of producing assistive technology for autistic children (52). The modules were developed through an iterative, user-centered design and evaluation process in conjunction with behavioral specialists and families with autistic children.

For example, Empowered Brain incorporates the Face2Face module, software that helps guide users to pay attention to socially salient visual stimuli (human faces; Figure 2 and Figure 3). The ability to pay attention to important social stimuli, and to direct gaze toward the most socially salient features of the face, has been identified as a key challenge in ASD [53].

**Figure 2.**
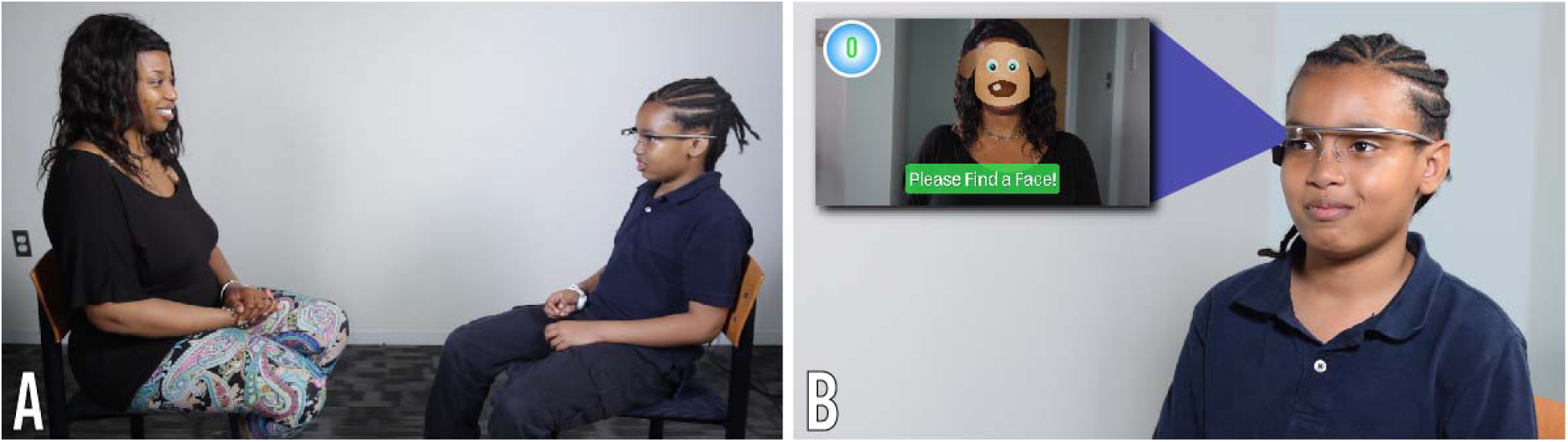
Demonstration of the use of the Empowered Brain System. (A) Child and partner sitting opposite one another with child wearing smartglasses. (B) Close-up view of the child wearing the smartglasses. The child can see the in-game view, displayed on the left side of the insert, through the optical display of the computerized smartglasses.

**Figure 3.**
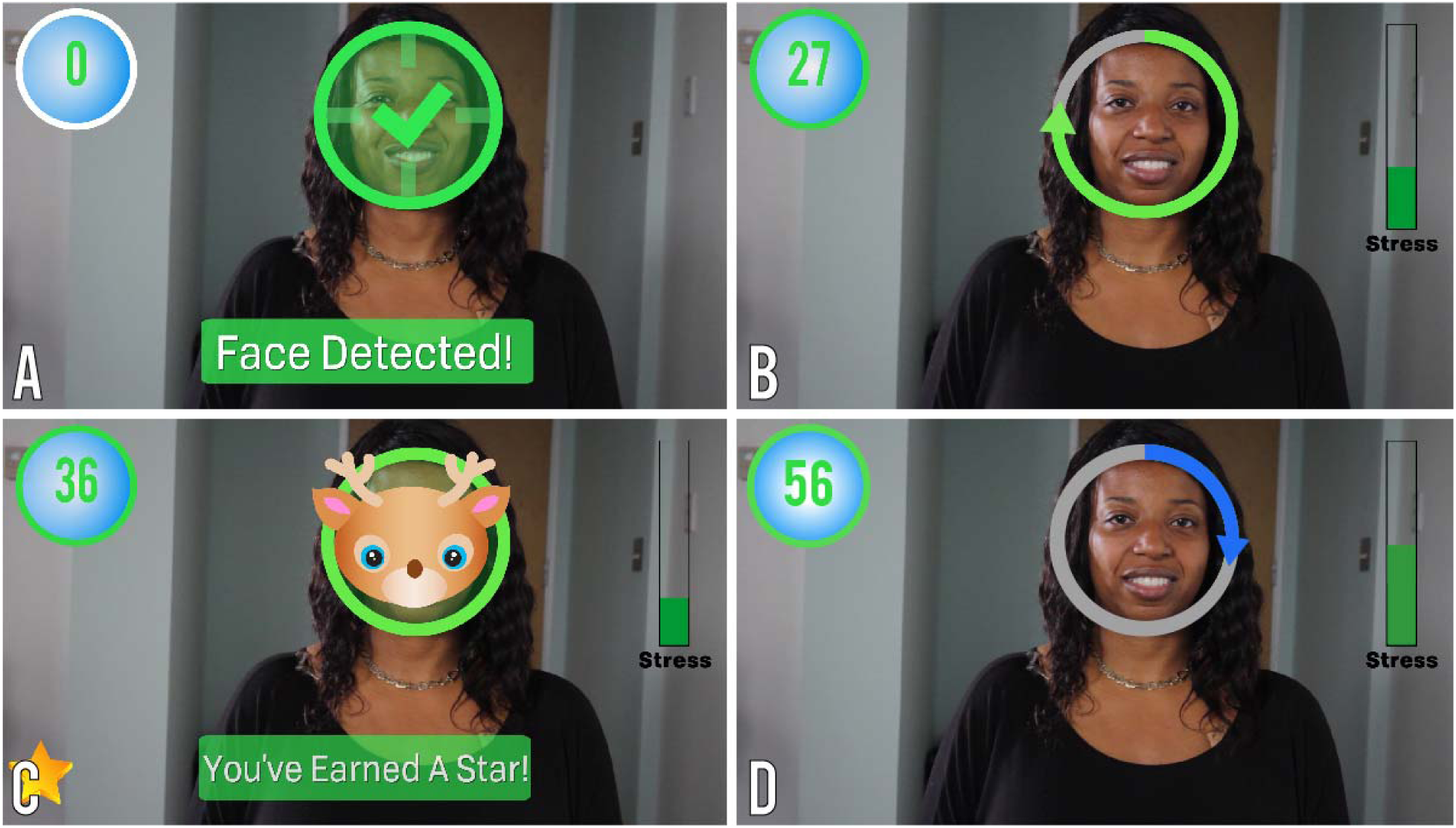
Empowered Brain’s Face2Face module. Face2Face is a two-player game that encourages face-directed gaze during social interactions.. (A) Child’s view on smartglasses: Face2Face detects the face in the field of view. (B) As the child maintains gaze towards the partner’s face, the ‘progress circle’ fills up and the child continues to earn points (upper left). (C) When the progress circle is full, the child earns a star (lower left) and a mask is displayed as a reward. (D) Another “progress circle” fills up as the child demonstrates continued gaze towards the partner’s face.

When the Face2Face module is running, Empowered Brain is able to identify the presence of human faces within its visual field and helps direct users toward the human faces through engaging visual cartoonlike images and guidance arrows. As the user pays more visual attention to the human face, they earn points and other in-game rewards. The points stop accumulating after a short period of time to avoid coaching users to stare. Figure 2 displays the relative positioning of the user and partner when using the Face2Face module.

Using a similar approach, Empowered Brain can detect not only human faces, but also human facial emotions when running the Emotion Charades module (Figure 4). In Emotion Charades, users have a gamelike experience in which they identify the emotions of another person. Empowered Brain rewards correct answers with in-game rewards or provides guidance when needed.

**Figure 4.**
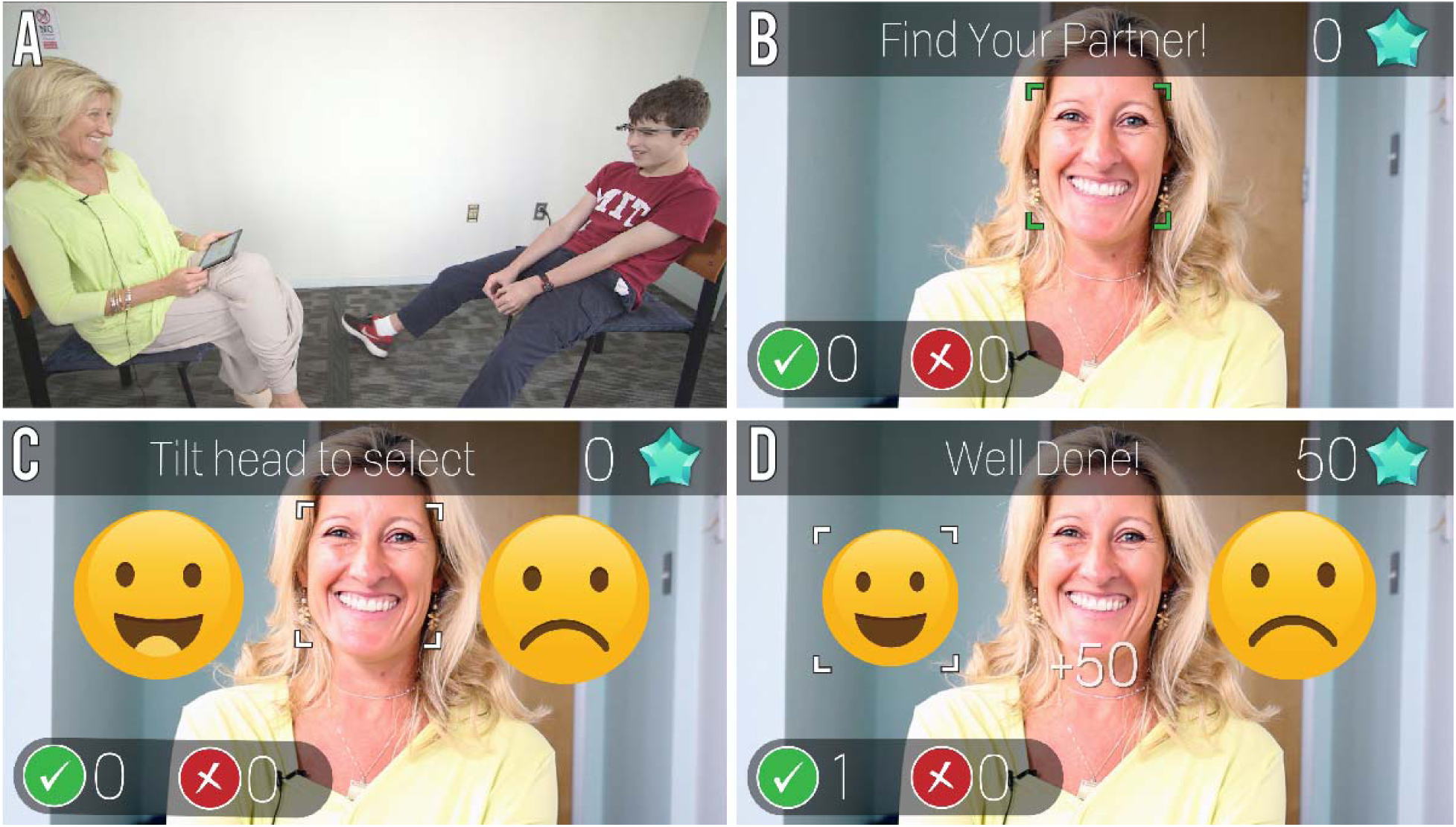
Empowered Brain’s Emotion Charades module. (A) Emotion Charades is a two-player game. The child, wearing smartglasses, sits across from his partner, whose app directs her to display an emotion. (B) Emotion Charades detects the partner’s face. (C) The partner displays an emotion (happy), which is detected by the system. The smartglasses shows the child two emotions. (D) The child tilts his head to select the emotion that matches the partner’s expression, earning points for correct responses (upper right).

Additionally, the system incorporates mechanisms to alter the difficulty associated with using each gamified app. One method is to alter the attentional challenge by displaying virtual elements that will either help to enhance attention or act as distractors to the social stimuli that the user is tasked to interact with. These virtual elements are overlaid over the user’s real-world view and include both dynamic real-time positional cues based on user movement and physiology, and reward-based virtual elements that provide feedback on the user’s in-app performance.

A series of research studies have investigated the use of Empowered Brain in ASD samples. The feasibility of using Empowered Brain in ASD was established during testing with 2 boys with ASD, both of whom demonstrated improvements in ASD symptoms as measured by the ABC [41]. Empowered Brain was safely used with no reported serious negative effects in 18 children and adults with ASD [54]. A separate report of 21 users with ASD found that Empowered Brain was well tolerated, with 91% of participants demonstrating tolerability across 3 separate measures [48]. The same study also found that 94% of participants reported the use of Empowered Brain to be comfortable. Additionally, the form factor of the computerized smartglasses of Empowered Brain has been described as desirable for school use by children with ASD [55]. Exploratory studies into the longitudinal use of Empowered Brain in school settings as a facilitative socio-emotional learning tool have been reported by multiple educators as having a positive impact on student learning and social communication [56]. Empowered Brain has also been found to improve social communication as measured by longitudinal educator and parental scores on the Social Responsiveness Scale, Second Edition, a reference standard validated measure of social communication functioning in ASD [57].

The facial affective analytics component of Empowered Brain was developed in partnership with Affectiva, an emotion artificial intelligence company. The Empowered Brain also uses experimental artificial intelligence technologies developed by Amazon. This work was also made possible by Google, Inc, now known as Alphabet, Inc, who provided substantial hardware and guidance in engineering. Brain Power, the company that developed Empowered Brain, has been a long-term Glass partner in the Glass Enterprise Partnership Program at X, a company of Alphabet, Inc.

## Methods

### IRB Approval

The methods and procedures of this study were approved by Asentral, Inc, Institutional Review Board (Newburyport, MA, USA), an affiliate of the Commonwealth of Massachusetts Department of Public Health.

### Participants

We recruited participants through a Web-based research interest form. Written consent was provided by the legal guardians of children and by cognitively abled adults. Participants between 7 and 17 years old provided written assent, when they were able to.

We questioned all caregivers of participants as to whether they had a history of ADHD and whether they were currently receiving treatment for ADHD. Additionally, all participants had a baseline ABC-H and Social Communication Questionnaire (SCQ) [58].

### Measures

The ABC-H is a subscale of the ABC and measures key ADHD symptoms such as inattention, impulsivity, and hyperactivity. The ABC has been extensively used in the developmentally disabled population, and the ABC-H has been used as an outcome measure of studies that have investigated the treatment of ADHD in populations with concurrent ASD [43-47]. In the assessment of ADHD symptoms in ASD, the ABC has shown itself to have superior psychometric properties, in particular validity [59] and reliability [60], compared with other rating scales of ADHD symptoms in children with developmental disorders (such as ASD). The ABC-H includes items that rate key ADHD symptoms. Specifically, the ABC-H assesses inattention and impulsivity through items that require the rater to assess whether their client is “easily distractible,” “does not pay attention to instructions,” “pays no attention when spoken to” or is “impulsive.”

Following these baseline assessments, we stratified the participants into high and low ADHD symptom groups based on their baseline ABC-H score. For the high ADHD group, we chose cutoff scores of 13 for male participants and 8 for female participants. These scores were determined by prior research that recorded the ABC scores for a sample of 666 people with developmental disorders, and found a mean ABC-H score of 13.38 for males and 8.12 for females [61].

Accordingly, we considered males with a score of 13 or higher and females with a score of 8 or higher to have high ADHD symptoms, and those with a lower score, to have low ADHD symptoms. Despite a history of ADHD diagnosis, we used the ABC-H as a stratification method, as it provided a numerical measure of recent (baseline) ADHD symptom burden. This numerical subscale allows for a more quantitative measure of change in rated items. While we obtained a clinical history of ADHD for the participants, the clinical diagnosis of ADHD in ASD is challenging [62], and it was only with the release of the *Diagnostic and Statistical Manual of Mental Disorders* (Fifth Edition) (DSM-5) [63] that it became possible to diagnose ADHD in an individual with ASD. Prior to the release of the DSM-5, the *Diagnostic and Statistical Manual of Mental Disorders* (Fourth Edition, Text Revision) specifically excluded a diagnosis of ADHD being made when an individual had a diagnosis of ASD. Therefore, while background information regarding ADHD history is important, we felt the ABC-H to be a more accurate measure of ADHD symptom load to determine the stratification into low and high ADHD symptom groups.

All participants had a baseline SCQ [58] as a validated measure of their ASD symptoms.

### Procedure

All participants were accompanied by a caregiver to the testing session. We oriented the participants and their caregivers to Empowered Brain and Google Glass and measured their ability to tolerate wearing the smartglasses. Once the participants showed they were able to wear the smartglasses for at least one minute, the participants were able to use Empowered Brain social communication modules and had a series of gamified experiences while interacting with their caregiver, including using Empowered Brain modules such as Face2Face and Emotion Charades (Figure 2, Figure 3). Empowered Brain modules help users to recognize and direct their attention toward socially salient stimuli such as human faces (in particular, the central part of the face, including eye regions), emotional facial expressions, and changes in the social environment. Participants and caregivers were able to verbalize any concerns or difficulties in using Empowered Brain both during and immediately after the session. We obtained an ABC-H score at 24 hours and at 48 hours after the session through the caregiver’s report. A clinically significant change in ABC-H was determined by a 25% or more change in the score, a standard that has previously been used in combination with another scale to determine responders to ADHD treatment in an ASD population [43].

Individuals who had expressed interest via the website signup but who had a known history of epilepsy or seizure disorder were not enrolled in this study. Users who had any uncontrolled or severe medical or mental health condition that would make participation in the study predictably hazardous were also not eligible for enrollment. We obtained information regarding potential exclusions to this study directly from the caregivers of the participants.

## Results

A total of 8 children, adolescents, and young adults with ASD signed up to take part in this research (average age 15 years, range 11.7-20.5; 7 male and 1 female participants). Table 1 and Table 2 summarize participant demographics. Half of the participants had a history of ADHD (n=4, 50%), 3 of whom were receiving active treatment at the time of testing. Of note, based on their ABC-H scores, 2 participants who had a previous diagnosis of ADHD were categorized in the low ADHD symptom group, while the remaining 2 were categorized into the high ADHD symptom group. The SCQ score demonstrated that participants represented a wide range of social communication abilities, from 11 to 28 points (mean score 18).

**Table 1.** Individual participant demographics (n=8).

| Participant identifier | Age (years) | Sex | SCQ | ADHD diagnosis | ADHD treatment |
| --- | --- | --- | --- | --- | --- |
|  | 12.6 | Male | 22 | No | No |
|  | 11.7 | Male | 11 | Yes | Yes |
|  | 12.4 | Male | 23 | No | No |
|  | 13.4 | Female | 18 | Yes | Yes |
|  | 16.9 | Male | 16 | Yes | No |
|  | 20.5 | Male | 28 | No | No |
|  | 19.4 | Male | 15 | Yes | Yes |
|  | 13.4 | Male | 12 | No | No |

<sup>a</sup>SCQ: Social Communication Questionnaire.
<sup>b</sup>ADHD: attention-deficit/hyperactivity disorder.

**Table 2.**
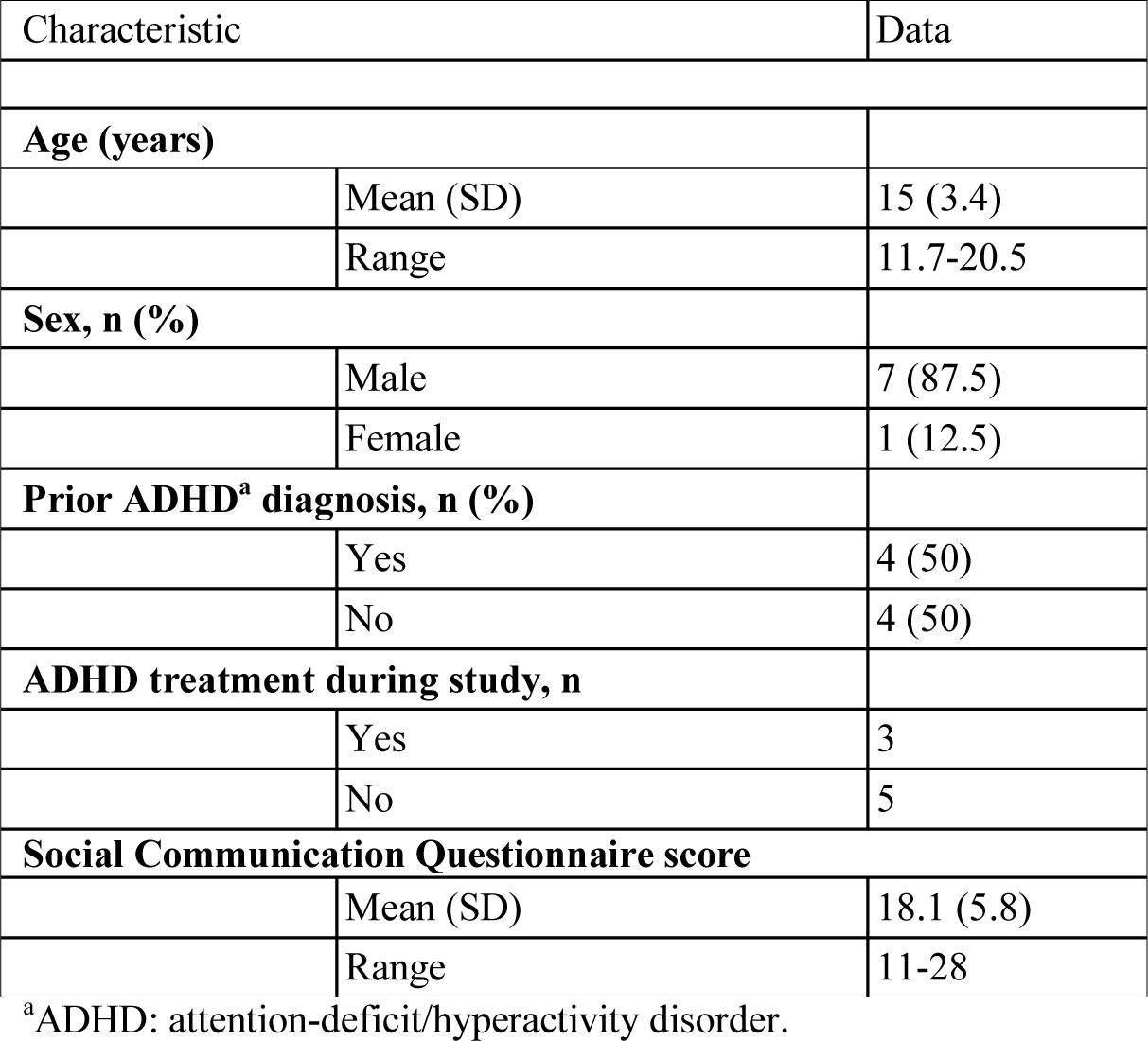
Overall participant demographics (n=8).

| Characteristic | Data |
| --- | --- |
| Age (years) |  |

|  |  |  |
| --- | --- | --- |
|  | Mean (SD) | 15 (3.4) |
|  | Range | 11.7-20.5 |
| **Sex, n (%)** |  |  |
|  | Male | 7 (87.5) |
|  | Female | 1 (12.5) |
| **Prior ADHD<sup>a</sup> diagnosis, n (%)** |  |  |
|  | Yes | 4 (50) |
|  | No | 4 (50) |
| **ADHD treatment during study, n** |  |  |
|  | Yes | 3 |
|  | No | 5 |
| **Social Communication Questionnaire score** |  |  |
|  | Mean (SD) | 18.1 (5.8) |
|  | Range | 11-28 |

<sup>a</sup>ADHD: attention-deficit/hyperactivity disorder.

All participants were able to use smartglasses and complete the coaching session. The high ADHD-related symptom group had a similar ASD severity to the low ADHD-related symptom group (SCQ score 18.5 vs 17.6, respectively), but consisted of younger participants (12.5 vs 17.6 years, respectively). Postintervention ABC-H scores were lower for most participants at 24 hours (n=6, 75%) and for all participants at 48 hours (n=8, 100%). At 24 hours after the session, average ABC-H scores decreased by 54.9% in the high ADHD symptom group and by 20.0% in the low ADHD symptom group. At 48 hours after the session, ABC-H scores compared with baseline decreased by 56.4% in the high ADHD symptom group (Table 3) and by 66.3% in the low ADHD symptom group (Table 4).

**Table 3.** High ADHDa-related symptom groups: ABC-Hb scores and percentage change relative to baseline (n=4).

| Participant identifier | Baseline score | 24-hour score | 24-hour % change | 48-hour score | 48-hour % change |
| --- | --- | --- | --- | --- | --- |
| 1 | 17 | 1 | -94.1 | 1 | -94.1 |
| 2 | 48 | 40 | -16.7 | 42 | -12.5 |
| 3 | 24 | 3 | -87.5 | 4 | -83.3 |
| 4 | 14 | 11 | -21.4 | 9 | -35.7 |
| Score mean (median) | 25.75 (20.5) | 13.75 (7) |  | 14 (6.5) |  |
| Mean Participant % change (median) |  |  | -54.9 (-52.4) |  | -56.4 (-57.5) |

<sup>a</sup>ADHD: attention-deficit/hyperactivity disorder.
<sup>b</sup>ABC-H: hyperactivity subscale of the Aberrant Behavioral Checklist.

**Table 4.** Low ADHDa-related symptom groups: ABC-Hb scores and percentage change relative to baseline (n=4).

| Participant identifier | Baseline score | 24-hour score | 24-hour % change | 48-hour score | 48-hour % change |
| --- | --- | --- | --- | --- | --- |
| 5 | 4 | 6 | 50 | 1 | -75 |
| 6 | 5 | 5 | 0 | 4 | -20 |
| 7 | 10 | 7 | -30 | 3 | -70 |
| 8 | 3 | 0 | -100 | 0 | -100 |
| Score mean (median) | 5.5 (4.5) | 4.5 (5.5) |  | 2 (2) |  |
| Mean Participant % Change (median) |  |  | -20 (-15) |  | -66.3 (-72.5) |

<sup>a</sup>ADHD: attention-deficit/hyperactivity disorder.
<sup>b</sup>ABC-H: hyperactivity subscale of the Aberrant Behavioral Checklist.

The high ADHD-related symptom group consisted of 4 participants who had an average ABC-H score of 25.75 at the start of the study (Table 3). The low ADHD-related symptom group consisted of 4 participants who had an average ABC-H score of 5.5 at the start of the study (Table 4). Figure 5 presents the ABC-H scores of the participants graphically.

**Figure 5:**
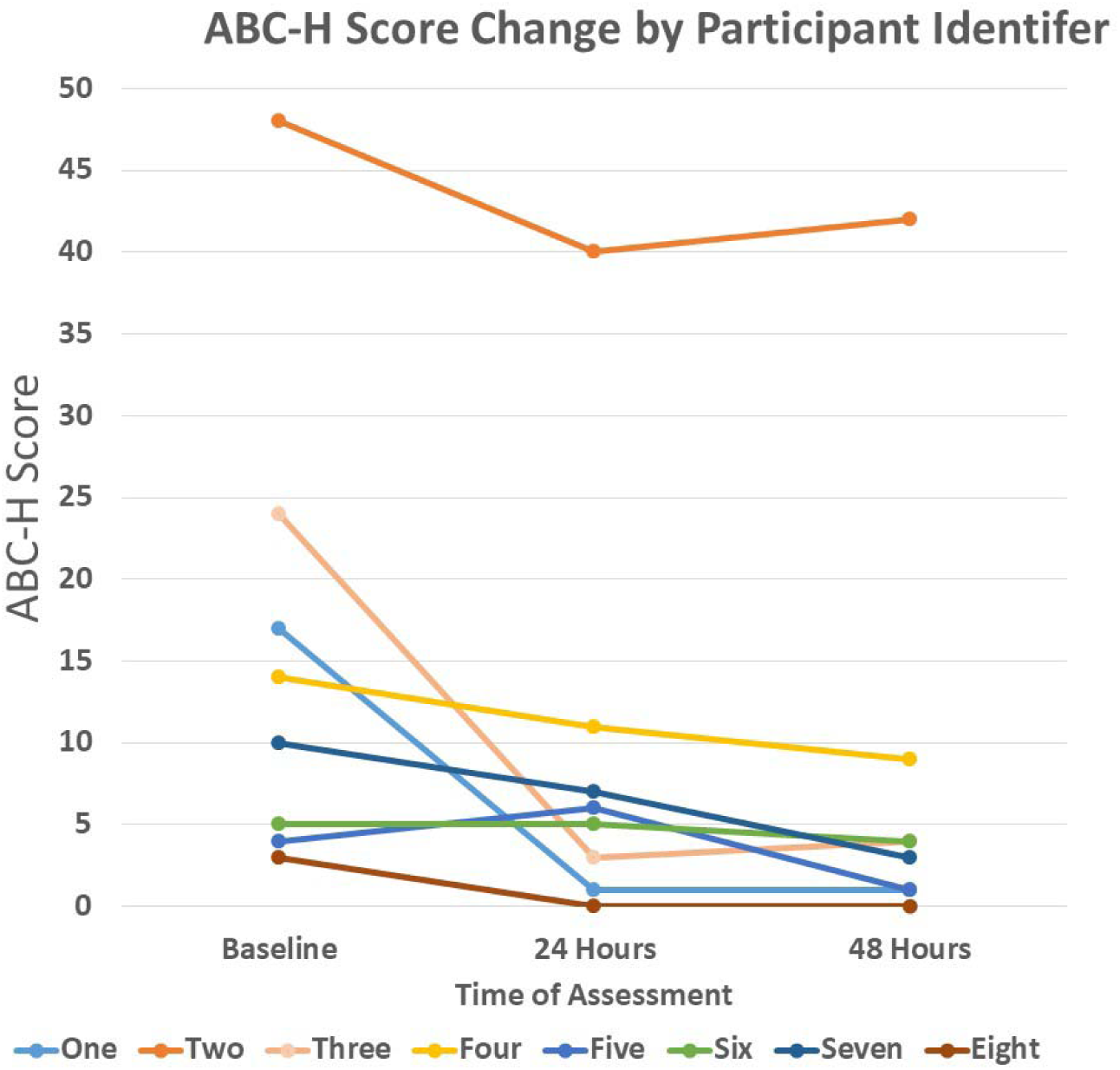
ABC-H score change by participant identifier from base to 48 hours post-intervention.

We noted that, while the participants in the high ADHD-related symptoms group were younger, they had a similar ASD symptom severity as measured by the SCQ to the low ADHD-related symptom group. It is perhaps not a surprise to find that the high symptom group was younger, given that the ABC-H subscale is weighted toward hyperactivity, and hyperactivity is an ADHD symptom that improves with age [64].

## Discussion

### Principal Findings

While many people with ASD struggle with symptoms of ADHD, including hyperactivity, impulsivity, and inattention, considerable gaps in knowledge remain in regard to their optimum assessment and management [3]. The combination of ADHD and ASD has been linked not only to greater impairment [10,11] and psychopathology [12,13], but also reduced access and response to psychopharmacologic treatment for ADHD [15,25]. These are important considerations given the substantial negative sequelae of untreated ADHD [29,30]. There has been increasing interest in developing digital interventions to address the symptoms of ASD or ADHD, but few technological interventions have been studied for people with ASD and ADHD symptoms. This population may potentially benefit from such approaches.

All participants in this study managed to complete the Empowered Brain intervention session without any reported negative effects, and all participants tolerated using smartglasses for the duration of the testing session. This is important, as a major limiting factor in the use of and continued engagement with assistive technologies is their usability, tolerability, and associated negative effects.

Most participants improved their ADHD-related symptoms following the intervention at both 24 hours (n=6, 75%) and 48 hours (n=8, 100%) after the session. We noted that 1 participant in the high symptom group had a greater reduction at 24 hours than at 48 hours, and that 1 participant in the low symptom group had an increase in ADHD-related symptoms at 24 hours followed by a large decrease from baseline at 48 hours.

Mean participant ABC-H percentage changes showed a significant response in the high ADHD-related symptom group (>25% improvement in ABC-H score) at both 24 (54.9% reduction) and 48 hours (56.4% reduction). The low ADHD-related symptom group appeared to show a response at 48 hours (66.3% reduction), but not at 24 hours (20% reduction). While the response of the low ADHD-related symptom group appeared to be of greater magnitude, we, who include subspecialist clinicians, advise caution in assessing the low ADHD-related symptom group, given the low baseline and small absolute score changes. Such results may render the findings in this group as not being as clinically significant as the outcomes associated with the high ADHD-related symptom group. An alternative explanation may also be put forward, and that is that the technology is especially impactful for individuals with a milder ADHD symptom burden. Future larger studies would allow for more robust statistical analysis and conclusions to be made.

### Limitations

There are several important limitations to this work that deserve mention. The number of participants in this pilot study was relatively small (N=8), although this is a sizeable sample relative to other research on novel technologies in ASD, especially in regard to other smartglasses research [41,48]. While quantitative approaches using validated scales are very useful, future research efforts would also benefit from the use of qualitative approaches, and some early results of such methods have been reported [56].

We should certainly consider the potential for an expectancy effect [65] in using this technology, especially given that the testing session was a novel experience for both the participant and the caregiver. However, the potential for this effect should also be tempered by our knowledge that transitions or new experiences have been associated with extreme distress in people with ASD, so much so that it is a characteristic part of diagnosis [63]. We noted that none of the participants encountered any noticeable distress or problems with using the smartglasses.

It would be useful for future research to incorporate a larger sample size, with more female participants. One can also see the benefit of age-matched neurotypical and ADHD-only controls. While the ABC-H is a very useful scale to use in this context, the use of broader ADHD-related measures would also provide for further insights. Despite our findings, the broader generalizability of our results to the wider ASD population will remain limited until further research is undertaken.

### Future Research

We hope that this study can pave the way for more funding and interest to study the potential application of this emerging technology to not just dually diagnosed ADHD and ASD samples, but more broadly to other categorical psychiatric diagnoses that are also associated with both attentional and social communication impairments. Such diagnoses include schizophrenia, bipolar disorder, and anxiety disorder. While deficits in social communication occur across many psychiatric disorders, most digital social communication interventions have been studied in ASD alone [66]. Therefore, research and development of an intervention that can target transdiagnostic symptoms could be valuable in both clinical scenarios and in research using the Research Domain Criteria framework.

Smartglasses may have a particularly unique role in this respect, as they also contain a variety of quantitative sensors that are located more closely than any other wearable technology to the main perceptual apparatus of the brain. These sensors can collect and help analyze sensor data to help classify human behavior. This digital phenotyping approach aims to detect changes in affective and behavioral states associated with both improvements and deteriorations in psychiatric disorders, and to help to identify new methods to subtype disorders [67].

Empowered Brain is being developed as a tool that can be used by caregivers, therapists, and educators to deliver socioemotional interventions to users with ASD. It is designed as a tool that can be used on a daily basis for 10 to 20 minutes per intervention. While this study reports on the impact of Empowered Brain on ADHD symptoms after a single intervention, further research is required to understand how the longitudinal use of this technology may affect ADHD symptoms in people with ASD. This technology also has the potential to address the attentional and social communication deficits that are transdiagnostically present across psychiatric disorders. Data collection via sensors that are proximally situated to the principal human perceptual organs may result in a more ecologically valid digital phenotyping approach. This digital phenotyping approach may aid the research of proposed developmental subtypes of a distinct ASD-ADHD combination disorder and, more broadly, psychiatric and brain injury-related disorders.

## Conclusion

Empowered Brain is a smartglasses-based social communication intervention that has been previously shown to improve social communication functioning in ASD. This study provides early evidence for the efficacy of Empowered Brain to reduce ADHD symptoms, such as hyperactivity, inattention, and impulsivity, in school-aged children, adolescents, and young adults with ASD. Our results also suggest that, despite concerns about increased distraction and reduced attentiveness, the Empowered Brain did not result in any increased ADHD symptoms in any of the participants at 48 hours after the intervention.

## Acknowledgments

The authors thank Google, Inc. for a generous grant and also the Glass team at X (formerly Google X) for technical guidance on Glass software development. This work was supported by the Office of the Assistant Secretary of Defense for Health Affairs through the Autism Research Program under Award No. W81XWH 17-1-0449. Early work to transform smartglasses into biomedical sensors was supported in part by the United States Army Medical Research and Materiel Command under Contract No. W81XWH-14-C-0007 (awarded to TIAX, LLC). Opinions, interpretations, conclusions, and recommendations are those of the authors and are not necessarily endorsed by the Department of Defense.

## Abbreviations

ABC-H: hyperactivity subscale of the Aberrant Behavioral Checklist
ADHD: attention-deficit/hyperactivity disorder ASD: autism spectrum disorder
DSM-5: Diagnostic and Statistical Manual of Mental Disorders (Fifth Edition)
SCQ: Social Communication Questionnaire

